# The cost of attentional reorienting on conscious visual perception: an MEG study

**DOI:** 10.1101/2020.12.05.413161

**Authors:** Alfredo Spagna, Dimitri J. Bayle, Zaira Romeo, Tal Seidel-Malkinson, Jianghao Liu, Lydia Yahia-Cherif, Ana B. Chica, Paolo Bartolomeo

## Abstract

How do attentional networks influence conscious perception? To answer this question, we used magnetoencephalography (MEG) in human participants, and assessed the effects of spatially nonpredictive or predictive supra-threshold peripheral cues on the conscious perception of near-threshold Gabors. Three main results emerged. (1) As compared with invalid cues, both nonpredictive and predictive valid cues increased conscious detection. Yet, only predictive cues shifted the response criterion towards a more liberal decision (i.e., willingness to report the presence of a target under conditions of greater perceptual uncertainty) and affected target contrast leading to 50% detections. (2) Conscious perception following valid predictive cues was associated to enhanced activity in frontoparietal networks. These responses were lateralized to the left hemisphere during attentional orienting, and to the right hemisphere during target processing. The involvement of frontoparietal networks occurred earlier in valid than in invalid trials, a possible neural marker of the cost of re-orienting attention. (3) When detected targets were preceded by invalid predictive cues, and thus reorienting to the target was required, neural responses occurred in left hemisphere temporo-occipital regions during attentional orienting, and in right hemisphere anterior insular and temporo-occipital regions during target processing. These results confirm and specify the role of frontoparietal networks in modulating conscious processing, and detail how invalid orienting of spatial attention disrupts conscious processing.

**Significance Statement:** Do we need to pay attention to external objects in order to become aware of them? Characterizing the spatiotemporal dynamics of attentional effects on visual perception is critical to understand how humans process and select relevant information. Participants detected near-threshold visual targets preceded by supra-threshold spatial cues with varying degrees of predictivity, while their brain activity was recorded using magnetoencephalography. Results demonstrated that valid predictive cues biased participants’ conscious perception through an early recruitment of frontoparietal regions, and that attentional costs associated to invalid predictive cues were related to activation of the right hemisphere ventral network. This work characterizes the neural dynamics associated with the cost of attentional reorienting on conscious processing.

## Introduction

The functioning of attention needs a balance between selecting relevant information for further processing, and ignoring irrelevant distractors. However, complete inhibition of distractors would not be adaptive, because the system needs to keep track of the novelty or potential danger of unattended information. In probabilistic environments, highly predictable events are associated with increased attentional selection of the relevant target and inhibition of the processing of distractors. However, in less predictable environments target selection and distractor inhibition might both be reduced.

Using the spatial orienting task (Chica et al 2014a, Posner 1980), in which a spatial cue precedes the relevant target, attentional selection and reorienting after distractor inhibition are measured as benefits and costs, respectively. As compared to a neutral condition in which the cue does not convey spatial information, benefits refer to faster reaction times (RTs) for targets presented at the valid location, while costs correspond to slower RTs for targets presented at the invalid location (e.g., Pestilli & Carrasco 2005). These costs are greater for spatially predictive peripheral cues than for nonpredictive ones (Chica et al 2013a), demonstrating an adaptive strategy in which the mental representation of the invalid location can be increasingly inhibited due to the infrequent appearance of targets at this location. In contrast, with spatially nonpredictive cues, the inhibition of the invalid location is reduced because targets are still likely to be presented at this location (Lasaponara et al 2011).

Manipulations of cue predictivity affect the speed of responses to valid and invalid locations, but also affect perceptual processes such as the conscious access of near-threshold information. Previous studies have demonstrated that spatially predictive cues are more effective in modulating conscious access than spatially nonpredictive cues (Chica et al 2011b, Dugué et al 2020). For near-threshold targets, while both cues made participants more liberal to respond to valid targets as compared to invalid ones, only predictive cues modulated perceptual sensitivity, which was increased for valid as compared to invalid trials (Chica et al 2011b).

Previous neuroimaging studies have demonstrated the importance of frontoparietal regions in supporting attentional functions. For example, in the model proposed by Corbetta and colleagues (Corbetta et al 2008, Corbetta & Shulman 2002), a dorsal frontoparietal network codes the relevant locations in space and helps selecting these locations during orienting. However, when relevant events are presented at unattended or unexpected locations, a more ventral frontoparietal network in the right hemisphere (including the right Temporo-Parietal Junction –TPJ, and the Inferior and Middle Frontal Gyri) allows attention to be re-oriented to the salient event. With supra-threshold targets, increasing cue predictivity is associated to reduced TPJ activity (Doricchi et al 2010, Shulman et al 2007). It was argued that during the cue period the TPJ builds-up and stores a template of attentional targets and when the target appears TPJ represents the target-to-template match or mismatch (Daffner et al 2003, Donchin & Coles 1988, Doricchi et al 2010, Kincade et al 2005, Knight & Scabini 1998, Serences et al 2005, Shulman et al 2007, Vossel et al 2006). This operation would help the monitoring and updating of probabilistic cue-target associations (Aston-Jones & Cohen 2005, Corbetta et al 2008, Donchin & Coles 1988, Lasaponara et al 2011).

The above-mentioned studies have explored the brain mechanisms associated to orienting of attention to supra-threshold stimuli; however, no study so far has compared the spatiotemporal dynamics of the brain mechanisms underlying attentional orienting using near-threshold targets preceded by spatially predictive and nonpredictive cues. Such an enterprise is important to explore the relationship between spatial orienting, expectation, and conscious perception. Here, we addressed two questions: (1) What are the spatiotemporal dynamics of the effects of peripheral cues on visual conscious processing? (2) How does cue predictivity modulate these effects? We recorded magnetoencephalography (MEG), capitalizing on its unique capacity to characterize a wide range of neural dynamics (Baillet 2017), while participants performed a version of the spatial orienting paradigm (Chica et al 2014a) with supra-threshold peripheral cues and near-threshold Gabor targets. In different experiments, spatial cues were either nonpredictive or predictive of the site of occurrence of targets. This setting enabled us to examine the effects of cues on conscious visual perception, both in terms of behavioral effects and of associated neural activity.

## Materials and Methods

### Participants

G*Power was used to estimate the sample size required to detect a difference in perceptual sensitivity for valid and invalid cue trials, we conducted a statistical power analysis using parameters from our previous study (Chica et al 2011b). For a t-test (match pairs), with alpha = 0.05, an expected power of 0.80, and an effect size of 0.81, the projected sample size needed was of n = 15 (two-tailed). We also conducted a statistical power analysis to estimate the sample size required to detect a difference in criterion for valid and invalid cue trials. For a t-test (match pairs), with alpha = 0.05, an expected power of 0.80, and the effect size of 1.53 as shown in previous research (Chica et al 2011b), the projected sample size needed was of n = 6 (two-tailed).

In total, 37 participants were recruited across two experiments. Eighteen participants completed the experiment with nonpredictive cues (age = 24 ± 3.13 years; age range = 22-33 years; 6M), and nineteen participants completed the experiment with predictive cues (age = 24 ± 3.79 years; age range = 20-32 years; 7M). Five participants had to be excluded from data analysis of the predictive cue experiment due to issues in the data quality of MEG recordings. All participants were right-handed, reported normal or corrected-to-normal vision, and gave written informed consent before participation. The two samples did not differ based on age (W = 154.50; *p* = 0.28; rank-biserial correlation = .23) or gender (W = 129.50; *p* = 0.91; rank-biserial correlation = .02). The study was promoted by the INSERM (C11-49) and approved by the Institutional Review Board of Ile de France I.

### Stimuli and Procedure

The experiments were compiled and run using E-Prime software (RRID: SCR_009567; Psychology Software Tools, Pittsburgh, PA) on a Windows XP desktop computer. All stimuli were presented on a gray background at the center of a black projection screen using a PROPixx projector (resolution: 1050 × 1400 pixels; refresh rate: 60 Hz) located outside the shielded recording room.

The display consisted of three black boxes (3.6° × 4.9° of visual angle) presented on a gray background; the central one was presented at the center of the screen and contained a fixation point (a black cross) at its center (see **Figure 1**). The other two boxes were located 6° of visual angle to the left and right side and 4° of visual angle below the central box, respectively, a setting created to maximize MEG responses from early visual areas (Portin et al 1999).

**Figure 1.**
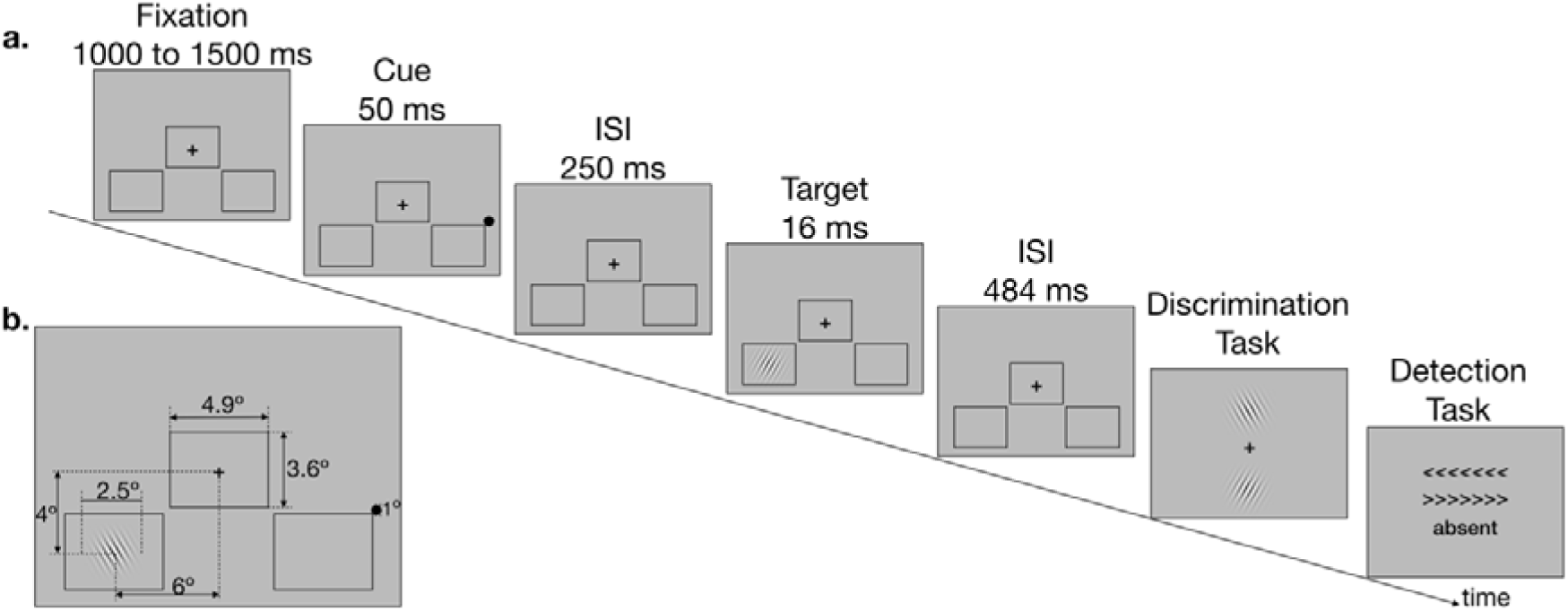
**a.** Schematic representation of the sequence of events in an invalid cue trial. **b.** Size of the stimuli and exact location of presentation on the screen. The experiment with nonpredictive cues (*NonPredCue*) and the experiment with predictive cues (*PredCue*) shared the same sequence of events. In the *NonPredCue* experiment 50% of cues were valid and 50% were invalid, while in the *PredCue* experiment 67% of cues were valid and 33% were invalid. ISI = inter-stimulus interval.

Participants sat in the MEG recording room, with the screen being positioned approximately 80 cm away from their eyes. Before the recording session began, participants were briefly instructed about the goal of the study and were then shown instructions on the screen. Each participant underwent a calibration session (mean duration, 6 min), during which target contrast (a Gabor patch) was manipulated to estimate the individual threshold for which the percentage of consciously perceived targets was ∼50%. The calibration block consisted of two randomly inter-leaved psychophysical staircases (one-up / one-down), theoretically converging toward a detection rate of 50%, while participants were engaged in the same paradigm as in the recording block (see below), except that the contrast of the stimuli was varied from trial to trial depending on their previous seen – unseen report in the corresponding staircase. Threshold contrasts were estimated separately for the valid and invalid cued locations to ensure a reasonable number of trials in the 4 experimental conditions of interest (seen-valid, seen-invalid, unseen-valid, unseen-invalid). The calibration block was followed by eight recording blocks (mean duration, 8 min per block).

During the recording blocks, each trial started with a fixation display, whose duration varied randomly between 1,000ms and 1,500ms. A cue was then presented, in the form of a black dot with a 1° diameter and presented for 50ms near the external upper corner of one of the two peripheral boxes. Three-hundred milliseconds after the presentation of the cue (an inter-stimulus interval - ISI - of 250ms), a Gabor patch (spatial frequency: 5 cycles per degree of visual angle; diameter: 2.5° of visual angle; orientation: chosen among 12 equally spaced between 0 and 180°, vertical and horizontal orientations being excluded) was presented as the target stimulus for 16ms in either the box to the left or to the right side of the display. After a 484ms ISI, participants performed two tasks sequentially: 1) a tilt discrimination task, which required participants to identify the orientation of the Gabor patch on a response box with 3 vertical buttons. By means of a button press, participants indicated the orientation of the grating among two possibilities presented vertically on the screen, distant by 3° from each other. Participants pressed the upper response button with their index finger to choose the upper orientation or the middle response button with their middle finger to choose the lower orientation. The location of the correct orientation was randomized. After the participants’ response to the tilt discrimination task, or after 3 s without response, a detection task, which required to press one of the three buttons of the response box to indicate whether the target was absent, or whether it had been presented in the left or right box. Two arrow-like stimuli (>>>>>> or <<<<<<) were presented above and below the fixation cross. Their respective position was randomized across trials. The word “absent” was presented under the arrow-like stimuli. When participants reported to have seen a stimulus, they pressed the upper or lower response button (with their index or middle finger, respectively) to indicate the visual hemifield of the target presentation. When participants reported not to have seen any stimulus, they pressed the lower response button with their ring finger. After the participants’ response, or after 3 s without response, the next trial began after a delay ranging from 1 to 1.5 s.

### Experimental designs and statistical analyses

In the nonpredictive cue (*NonPredCue*) experiment, each of the eight MEG recording blocks consisted of 110 trials, including 88 stimulus-present trials (in which stimuli at threshold contrast were presented either in the left or right lower visual quadrants) and 22 stimulus-absent trials (in which no stimulus was presented). The total number of trials was 880, with 50% valid cue trials (352 trials), 50% invalid cue trials (352 trials), and 176 catch trials (i.e., trials in which no target was presented). Half of the targets were presented at the cued location (valid cue condition); the other half was presented at the uncued location (invalid cue condition). Trials within a recording block were presented in a different randomized order for each subject.

In the predictive cue (*PredCue*) experiment, parameters of stimulus size and timing of presentation were the same as those used in the nonpredictive cue experiment, except that the total number of trials on each of the eight MEG recording blocks consisted of 784 trials, with 67% valid cue trials (448 trials), 33% invalid cue trials (224 trials), and 112 catch trials. Participants were instructed about the cue predictivity before beginning the session. Notably, previous work has shown that effects of cue predictiveness could be observed even when participants are not made aware of it (Bartolomeo et al 2001, Bartolomeo et al 2008, Decaix et al 2002, López-Ramón et al 2011).

### MEG recordings

Continuous MEG recordings were conducted at the CENIR (http://www.cenir.org) with an ELEKTA Neuromag TRIUX^®^ machine (204 planar gradiometers and 102 magnetometers) located in a magnetically shielded room with a sampling frequency rate of 1kHz and a bandwidth ranging from of 0.01 to 300 Hz. The recordings were then MaxFiltered (v2.2) (Taulu & Simola 2006) to attenuate environmental noise, Signal Space Separation (SSS) was then implemented, automatic detection of bad channels was conducted, data were filtered (1 to 250 Hz), and resampled at a rate of 250Hz, and then converted in the Fieldtrip structure (RRID: SCR_004849; http://www.fieldtriptoolbox.org/) (Oostenveld et al 2011) to conduct further preprocessing and analytic steps. Cardiac activity (electrocardiogram – ECG), vertical and horizontal Electroculogram (EOG) signals were also recorded together with the electrophysiological data. The exact timing of the presentation of the stimuli onset was corrected in accordance to the signal received from a photodiode located in the MEG room, in order to adjust to the delay produced by the refresh rate of the projector.

### Preprocessing and Artifact Rejection

Additional preprocessing steps were conducted using Fieldtrip and included an initial visual inspection of the recordings conducted by two of the authors (D.J.B. and Z.R.) to exclude segments with artifacts and ensure data quality control. EOG recordings from both vertical and horizontal sensors were then used to reject trials in which eye movements (beyond 3°) occurred. Rejection thresholds for both horizontal and vertical EOG traces was set to ± .66V, corresponding to a deviation greater than 3° of visual angle (and with the target at 6° of visual angle). Trials with excessive eye movements and eye blinks (∼10.52% of trials) were rejected offline from the MEG traces according to the 3° threshold mentioned above. Signal from the photodiode was used to discard 1) trials with a delay between the trigger and the photodiode greater than 300ms; 2) trials with a delay between the cue and the target greater than 827ms; 3) trials in which the delay between the trigger of the cue and the photodiode was greater than 40ms or smaller than 30ms, for a total of ∼1% of the trials. Last, trials contaminated by muscular activity (jump or movement) were rejected manually upon visual inspection (∼15%).

For the experiment with nonpredictive cues, out of the 15,840 trials acquired, 8,517 trials were analyzed (*right visual field*: seen invalid = 1,033; seen valid = 1,284; unseen invalid = 1,056; unseen valid = 882; *left visual field*: seen invalid = 1,191; seen valid 1,176; unseen invalid = 956; unseen valid = 939).

For the experiment with predictive cues, out of the 10,796 trials acquired in total, 7,454 trials were analyzed (*right visual field*: seen invalid = 396; seen valid = 1,588; unseen invalid = 812; unseen valid = 920; *left visual field*: seen invalid = 639; seen valid = 1,144; unseen invalid = 576; unseen valid = 1,379) (see **Table S1** for a subject-by-subject breakdown of the number of trials in each condition remaining after artifact rejection).

This procedure led to a difference of about 4% in the number of consciously reported trials that underwent MEG analyses between the *PredCue* (3,767 /10,796 = 34%) and *NonPredCue* (4,686 /15,840 = 30%). We did not anticipate this difference to have an influence on the contrasts of interest for this study, as further elaborated in the discussion. Supplementary Table (S1) shows that after discarding trials we were still able to keep a great part of the *predictivity* difference across the two tasks, and that more than 400 trials total were analyzed per each participant.

### Event-Related Magnetic Fields

Data from 102 NEUROMAG channels was analyzed in this study. A Matlab® script was used to separate the MEG continuous recordings into 2300ms- long epochs (ranging from -1000 before the cue and 1300ms after the cue), and epochs from the eight experimental conditions from each participant were then imported into Brainstorm (Tadel et al 2011). For each condition, event-related magnetic fields were then averaged (weighted) along their entire length (2300ms).

### Source reconstruction

Signal amplitude from the 15,002 cortical elemental dipoles underlying the signals measured by the sensors were then estimated from the epochs using the weighted minimum norm estimation (wMNE) imaging method as implemented in Brainstorm (Tadel et al 2011). The number of dipoles used in our study is a standard in MEG literature, and is meant to cover the entire cortical surface and its folded anatomy (Tadel et al 2019). The wMNE method identifies a current source density image fitting the data through the forward model, and then favors solutions that are of minimum energy by using source covariance as a prior. wMNE modeling of the sources was used because it is considered to be the gold standard in the field, because it balances between a conservative approach in reconstructing sources of MEG signal, with reliability of statistical inference across participants, and being computationally efficient (less resource expensive)(Baillet et al 2001, Tadel et al 2019). A noise covariance matrix was estimated for each subject from the recordings using the pre-stimulus interval (-1,000 to -2ms before the presentation of the cue). Estimating the baseline covariance (i.e, noise covariance) over such an amount of time (approximating one second) allows us to assume a sufficient regularization term of the solution (noise from brain activity at rest though to be unrelated to our task conditions). Constrained source covariance model was used to model one dipole, oriented normally to the surface, a choice that allows us to study the cortical regions associated with the interaction between attention and consciousness. Further processing conducted on the sources per participant consisted of z-score transformation of the signal with reference to the baseline (from -1,000 to -2ms). A default value was chosen for the spatial smoothing kernel (FWHM = 3mm), and was then applied on the sources that were then re-interpolated (projected) on a common template (MNI 152).

### MRI recordings

High-resolution T1-weighted structural MRI images (MPRAGE sequence, flip-angle, 9; Repetition Time, 2300ms; Echo Time, 4.18ms; voxel size: 1 × 1 ×1 mm) were acquired for each participant using a 3-T Siemens, TRIO whole-body MRI scanner (Siemens Medical Solutions, Erlangen, Germany) located at the CENIR MRI center (Salpetriére Hospital, Paris, France). After acquisition, images were then segmented using the FreeSurfer “recon-all” pipeline (Fischl 2012), and imported in Brainstorm (Tadel et al 2011) for co-registration purposes. MEG sensors and structural MRI images were first manually aligned using the nasion/left ear/right ear (NAS/LPA/RPA) fiducial points recorded in the MEG file and in the MRI MNI coordinates. Co-registration was then further refined using the “refine using head points” option on Brainstorm, which uses an iterative closest point algorithm to fit the head shape and the digitized scalp points. Additional details about the MRI-MEG co-registration steps as done in Brainstorm can be found here (Tadel et al 2019).

### Behavioral Data Analysis

Even though target contrast was adjusted to detect ∼50% of the targets presented at the valid and invalid location during calibration, we used the signal detection theory (SDT) to estimate changes in the signal to noise ratio (*a’*) as a function of cue validity condition, and to investigate whether the presence of a cue could bias the observer towards a more liberal or conservative reporting threshold (*beta’’*). A more liberal criterion indicates that a participant is more likely to indicate the presence of a target even under conditions of increased perceptual uncertainty (i.e., at a lower level of visual stimulation received), and therefore also more likely to report false alarms. On the other hand, a more conservative criterion indicates that a participant will report a target under conditions of higher visual stimulation, a strategy that may result in an increase of missed targets. We chose to use nonparametric measures of perceptual sensitivity and response criterion because some participants made no false alarms (Pollack & Norman 1964). We computed the mean percentage of seen targets when the Gabor was presented (hits) and when the Gabor was absent (false alarms; FA). The following formulas were used to calculate a’ and *beta’’*:

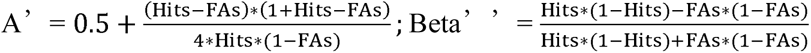

A’ values usually range between 0.5 (the signal cannot be distinguished from the noise) and 1 (perfect performance). For Beta”, values close to 1 indicate a conservative criterion, whereas values close to − 1 indicate a liberal criterion (Stanislaw & Todorov, 1999). Both a’ and beta’’ were estimated separately for valid and invalid trials. Paired sample t-tests were used to assess differences in either index between valid and invalid trials. Wilcoxon signed-rank test was used to assess potential differences in the thresholds sampled during the calibration blocks separately for valid and invalid trials.

A 2 (*NonPredCue*, *PredCue*) x 2 (valid, invalid) repeated measures ANOVA was also conducted on hits, a’, and beta’’ to assess cross-experiments differences. Further, a 2 (seen, unseen) x 2 (valid, invalid) repeated measures ANOVA was conducted on the percentage of correctly discriminated responses (i.e., accurate response in the discrimination task). Analyses on RTs, were considered not informative due to the experiments’ procedure, and are not reported (see **Supplementary Material**).

### MEG data analysis

To examine whether neuronal activity induced by visual cues can enhance conscious perception, permutation *t*-tests with spatiotemporal clustering (Maris & Oostenveld 2007) were conducted separately for the *NonPredCue* and *PredCue* experiments. The formula used to capture the interaction was: [valid (seen *minus* unseen) *minus* invalid (seen *minus* unseen)]. A positive *t* value indicates an increased awareness effects for valid as compared to invalid trials. Brain activation was examined in the time window between 0 and 800ms (locked to cue onset), with the number of permutations set to 1,000 and the alpha threshold level set to 0.05 for all tests.

To examine whether predictive and nonpredictive visual cues are associated with distinct brain dynamics, spatiotemporal cluster-based permutation t-tests were also conducted to compare cross-experiments differences in the activation of brain regions, separately for valid and invalid trials.

### Control Analyses

(1) A 2 (seen, unseen) × 2 (left visual field, right visual field) × 2 (valid, invalid) repeated measures ANOVA was conducted separately on the percentage of trials analyzed in the MEG analyses to identify potential differences across task conditions. (2) Only 2 subjects participated in both experiments. To ensure that cross-experiments differences were not due to the different composition of the samples, ANOVAs on MEG data were repeated for these two subjects separately. Results on this sub-sample showed comparable results than those obtained from the group analyses, supporting the analytic approach conducted in this study.

## Results

### Do predictive cues affect behavioral responses to near-threshold Gabors?

Participants detected more targets in valid trials than in invalid trials (mean ± SD valid: 53.33% ± 7.30; invalid: 42.60% ± 5.60; *t*(13) = -7.57; *p* <.001), and adopted a more liberal criterion after valid trials than for invalid trials (mean ± SD valid: 0.56 ± 0.34; invalid: 0.88 ± 0.15; *t*(13) = 3.99; *p* <.05) (see **Fig 2a** and **2b**). The distributions of the thresholds established during calibration separately for valid and invalid cue trials resulted statistically different (Wilcoxon signed-rank test: W = 75.50; *p* <.05). Yet, perceptual sensitivity (*a’*) was comparable for valid trials (0.81 ± 0.08) and invalid trials (0.84 ± 0.03; *t*(13) = 1.56; *p* =.18).

**Figure 2.**
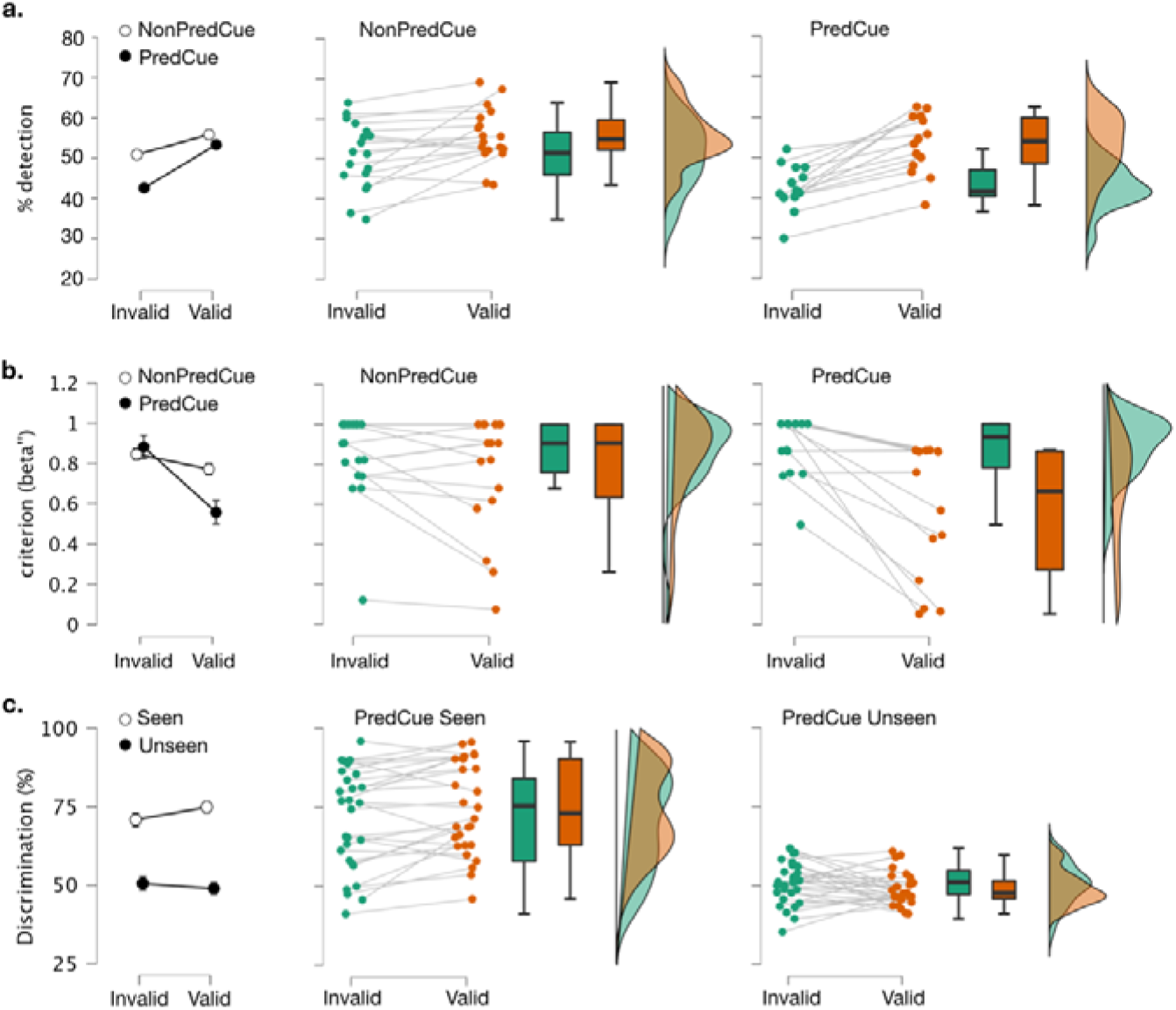
**a.** percentage of detected targets on the *NonPredCue* and *PredCue* experiments; **b.** criterion on both the *NonPredCue* and *PredCue* experiments; **c.** percentage of discriminated targets for the *PredCue* experiment for the seen (left panel) and unseen (right panel) trials.

Discrimination accuracy was also greater for *seen* trials (72.87% ± 15.21%) than for *unseen* trials (49.71% ± 5.95%) (main effect of *Awareness*: F_(1,13)_ = 31.67; *p* < .0001; η^2^ = .71; see **Fig 2c**). The significant interaction between *Awareness* and *Validity* (F_(1,13)_ = 11.9; *p* < .01; η^2^ = .48) showed that the valid / invalid difference was significant for seen trials (valid: 74.90% ± 14.21; invalid: 70.86% ± 15.70; *p* < .01), but not for unseen trials (valid: 48.91% ± 3.48; invalid 50.51% ± 3.88; *p* = .19). The interaction between *Visual Field* and *Validity* (F_(1,13)_ = 8.89; *p* < .05; η^2^ = .41) showed that participants discriminated more validly cued targets than invalidly cued targets in the right visual field (valid: 64.24% ± 9.92, invalid: 60.08% ± 12.26, *p* < .01), but not in the left visual field (valid: 59.56% ± 9.62; invalid: 61.29% ± 10.04 *p* = .21). No other interaction reached significance [*Awareness* x *Visual Field* and (F_(1,13)_ = 1.66; *p* = .22; η^2^ = .11); *3-way* interaction (F_(1,13)_ = 0.02; *p* = .88; η^2^ = .01)].

### Do predictive cues affect neural responses associated with the conscious report of Gabors?

We addressed this question by examining brain responses to the interaction term [valid (seen *minus* unseen) *minus* invalid (seen *minus* unseen)]. Two clusters exceeding the threshold of randomization distribution under H0 emerged (*ps < .05*), the first lateralized to the left hemisphere and observed during the cue-target period in temporo-parietal regions; the second, lateralized to the right hemisphere and occurring during the post-target period in the frontal-eye fields, inferior parietal and temporo-parietal regions, and then in the temporal lobe.

Control analyses showed that the percentage of trials analyzed in the left or right visual field did not differ as a function of cue validity (*p* = .84). There was an interaction with the factor awareness (*p* < .05). However, post-hoc comparisons (Bonferroni corrected) did not reveal any significant difference across any of the comparisons.

To further characterize the significant spatiotemporal pattern associated with the interaction, we analyzed the awareness effect (seen *minus* unseen) separately for valid and invalid predictive cues (**Figure 4**). For valid trials, we observed two clusters, both during the post-target period: the first lateralized to the right hemisphere and beginning in the occipito-temporal regions (around 400ms post-target) and then reaching into parietal (450ms) and wide-spread frontal (480ms) areas. The second cluster, lateralized to the left hemisphere, was observed in the occipital region (around 410ms), but reached earlier into parietal (420ms), insular (around 440ms) and inferior frontal areas (470ms), as compared with the first cluster.

**Figure 3.**
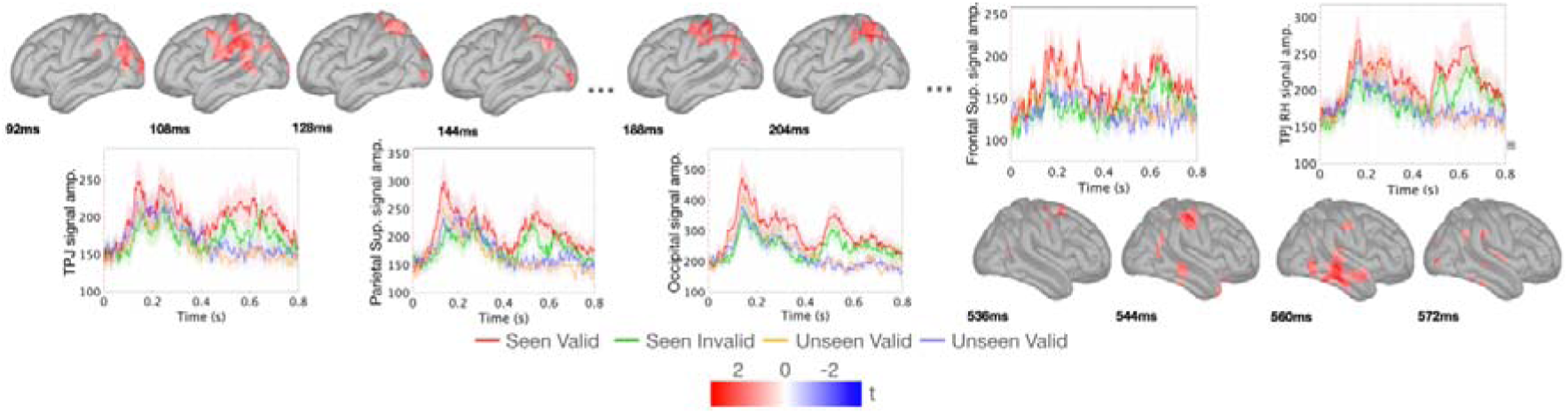
When preceded by valid predictive cues, seen targets evoked increased responses in two clusters of brain activity, as compared to invalidly cued targets. Cluster 1 occurred in the 90-210ms time window after cue onset and was lateralized to the left hemisphere; Cluster 2 occurred in 530ms time-window after cue onset and was lateralized to the right hemisphere.

**Figure 4.**
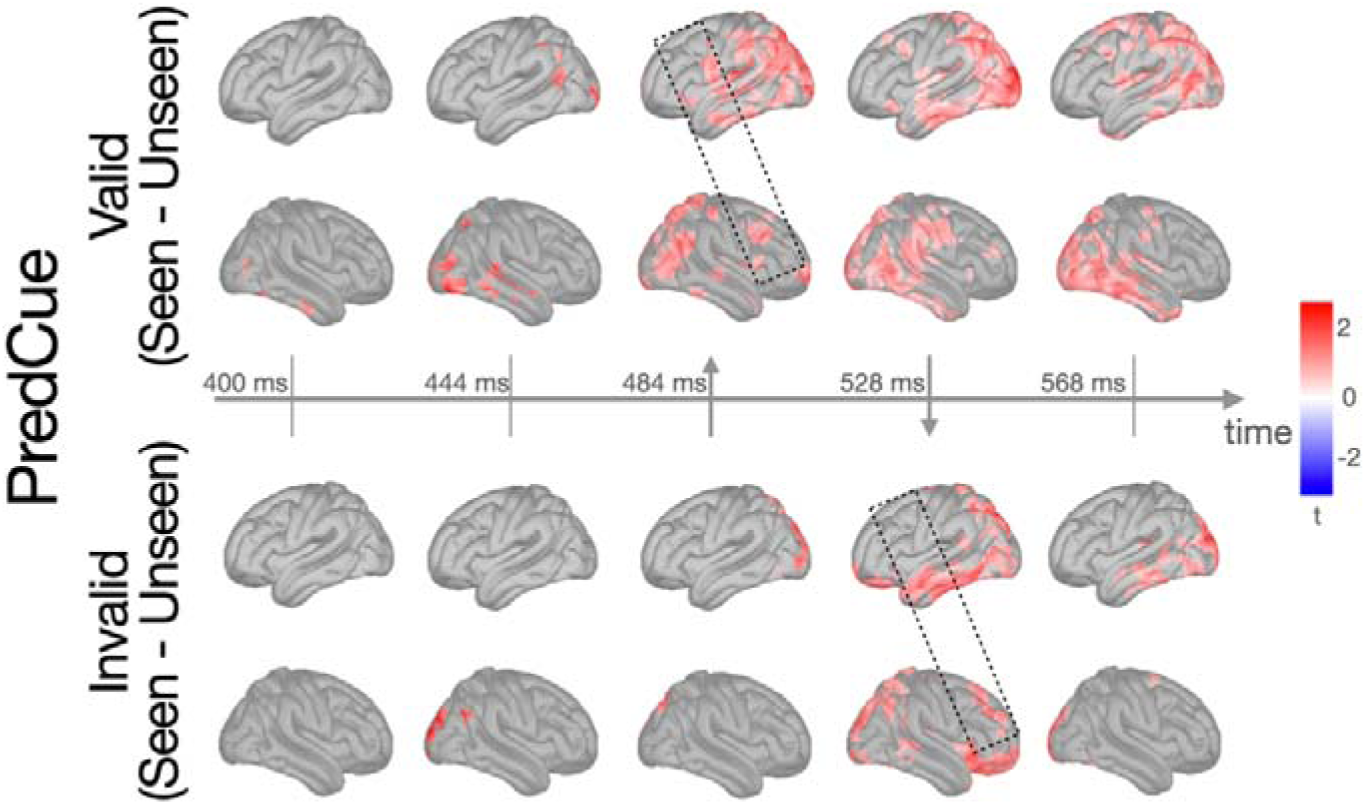
Analysis of the Awareness effect observed in the *PredCue* experiment analyzed separately for valid and invalid cue trials. Temporal dynamics of the awareness effect seemed to differ between valid and invalid trials, with prefrontal activations (indicated by rectangular boxes) occurring earlier in valid trials.

For invalid trials, we observed two post-target clusters: the first lateralized to the right hemisphere and beginning in the occipital region (around 410ms) and including also superior parietal areas at around 470ms. Activity in a cluster located in the right TPJ and in the ventromedial prefrontal cortex (around 490ms) preceded that of lateral frontal regions (middle, 504ms; superior including FEF, around 510ms) before spreading to fronto-parieto-occipital activation. The second cluster, lateralized to the left hemisphere was temporally delayed compared to the contralateral activation. It began in the occipital lobe around 440ms, and then spread to superior parietal regions at 470ms; only later did it involve middle and inferior temporal regions, TPJ, and ventromedial prefrontal cortex at around 520ms.

### Do nonpredictive cues affect behavioral responses to near-threshold Gabors?

Participants detected more targets in valid trials than in invalid trials (mean ± SD valid: 55.80% ± 6.90; invalid: 50.90% ± 8.30; *t*(17) = -3.25; *p* <.01). No significant differences occurred between valid and invalid trials in response criterion (mean ± SD valid: 0.77 ± 0.29; invalid: 0.85 ± 0.22; *t*(17) = 1.84; *p*=.084), perceptual sensitivity (*a’*) (mean ± SD valid: 0.86 ± 0.06; invalid: 0.86 ± 0.04; *t*(17) <1), or perceptual thresholds resulting from the calibration for valid trials (43.98%) and for invalid trials (44.00%; Wilcoxon signed-rank test: W = 82.00*; p =* .90).

As expected, participants were more accurate in discriminating the orientation of *seen* targets (79.2% ± 14.0) than that of *unseen* targets (50.2% ± 7.0) (main effect of *Awareness*: F_(1,17)_ = 85.60; *p* < .0001; η^2^ = .84; see **Figure 2)**. No other factors or interactions reached statistical significance.

### Do nonpredictive cues affect neural responses associated with the conscious report of near-threshold Gabors?

We addressed this question by examining brain responses to the interaction term [valid (seen *minus* unseen) *minus* invalid (seen *minus* unseen)]. No cluster exceeding the threshold of randomization distribution under H0 emerged (*ps < .05*)(see **Supplementary Material**). Control analyses showed that the percentage of trials analyzed in the left or right visual field did not differ as a function of either awareness (*p* = .51), or cue validity (*p* = .10).

### Cross-experiment comparisons: different effects of nonpredictive and predictive cues on visual conscious perception

We observed a significant interaction between *Validity* and *Experiment* on both the percentage of detected targets (F_(1,30)_ = 7.25; *p* < .05; η^2^ = .19) and the response criterion (F_(1,30)_ = 8.61; *p* < .01; η^2^ = .22). When targets were presented at invalidly cued locations, the percentage of correct detections were greater in the *NonPredCue* experiment (51% ± 0.8) than in the *PredCue* experiment (43 ± 6%; t = 3.24; *p_Bonf_* < .05), while the percentage of detected targets was comparable between the experiments for valid trials (*NonPredCue*: 56% ± 7; *PredCue* 53% ± 7; *p_Bonf_ ∼* 1). Concerning criterion, planned comparison between experiments for valid and invalid trials did not reach significance (both *p*s > 0.152).

For the discrimination task, there was an interaction among the factors of *Experiment* and *Validity* (F_(1,30)_ = 5.35; *p* < .05; η^2^ = .15). However, none of the pairwise comparisons reached statistical significance. Average accuracy did not differ between the two tasks (F_(1,30)_ = 1.88; *p* = 0.18; η^2^ = .06).

Results of the ANOVA conducted on the MEG data revealed three clusters of activation that reached statistical significance for the invalid trials, while no significant result was obtained for valid trials (see **Figure 5**). The first cluster appeared during the cue-target period and was lateralized to the left hemisphere; it started in the anterior portion of the inferior temporal lobe (around 250ms) and then reached into the insular cortex (276ms), TPJ and occipito-parietal junction (288ms), where it remained active until 320ms. The second and third clusters appeared both during the post-target period and were lateralized to the right hemisphere beginning in the inferior temporal lobe (570ms), TPJ and anterior temporal lobe (576ms), the middle insular cortex (604ms), a widespread activation occipito-temporal regions (around 650ms) especially along the medial surface, the ventromedial prefrontal cortex, and remained active until 670ms.

**Figure 5.**
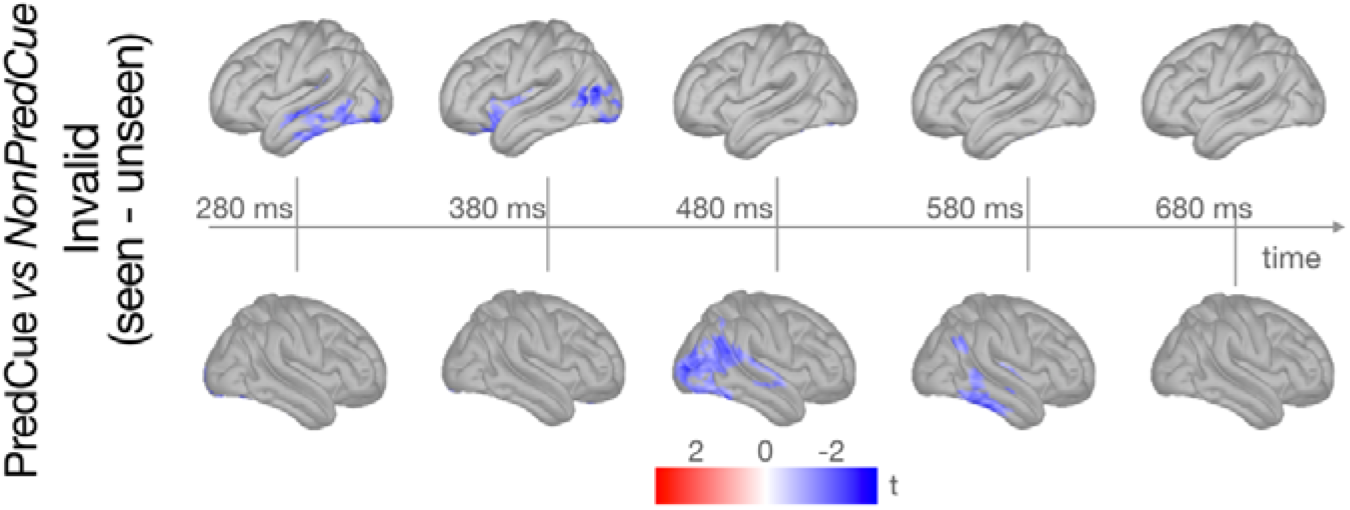
Cross-experiment analyses of neural data. We observe a concordant pattern in the results from analyses on the percentage of correctly detected targets and from the MEG data, with invalid trials in the *NonPredCue* experiment characterized by higher activation compared to those in the *PredCue* experiment.

## Discussion

Our results elucidate the patterns of interaction between spatial attention and consciousness and underlying spatiotemporal dynamics, and how they vary as a function of cue predictivity. Altogether, the contributions of nonpredictive and predictive valid cues in improving target detection indicate that when attentional resources are allocated towards near-threshold visual information conscious access is improved, and that the brain seems to exploit the predictivity of the cue at the expense of increasing costs associated to attentional reorienting.

### Behavioral effects of spatial cues on conscious reports

Behavioral results align with previous evidence in revealing a pattern of interaction between attention and consciousness (Ling and Carrasco, 2006; Liu et al., 2009; Chica et al., 2011b; Sergent et al., 2013; Botta et al., 2017; Vernet et al., 2019). First, both predictive and nonpredictive cues modulated the proportion of detected targets, but only spatially predictive cues significantly modulated response criterion; they produced a greater cost to detect targets presented at invalid locations. Further, conscious reports decreased after invalid trials, especially when cues were spatially predictive, and therefore invalid trials were infrequent. Rather than improving conscious perception at the valid location, making the cues spatially predictive increased attentional costs to reorient attention to invalid locations, as previously shown in studies using fMRI (Doricchi et al 2010) and EEG (Lasaponara et al 2011) with supra-threshold cues and targets.

### Increased frontoparietal activation in predictive valid compared to invalid cues

The interaction between attentional orienting and conscious awareness observed at the behavioral level in the experiment with predictive cues was accompanied by increased frontoparietal activation in valid compared to invalid trials. Our results are consistent with longstanding evidence regarding the role of the frontoparietal network in supporting attentional mechanisms (Corbetta and Shulman, 2002; Fan et al., 2005; Chica et al., 2013b; Xuan et al., 2016), and expand on the role of this network in enhancing subjects’ visual awareness. Yet, thanks to the spatiotemporal resolution of MEG, we were able to pinpoint the temporal differences in the recruitment of this neural network between valid and invalid cues. Peripheral predictive cues produce strong expectation regarding the location where the target will be presented, an effect that is beneficial for valid trials, but detrimental in the less frequent cases when the cue is invalid. Our follow-up analyses showed a delay in the recruitment of frontoparietal regions for reported invalid targets, as compared to unreported targets. Such delay is likely to be a neural marker of the attentional cost of reorienting attention, caused by being misled to attend to the incorrect location (Corbetta et al 2008). Contrary to recent claims (Raccah et al 2021), our interpretation reinforces the central role of frontal and prefrontal regions in conscious visual processing (Chica et al 2014b), a claim that is also supported by the absence of such effects in the experiment with nonpredictive cues, which did not induce any probabilistic expectations.

### Increased temporo-occipital activation for nonpredictive compared to predictive invalid cues

Cross-experiment comparisons further elucidated the neural pattern associated with manipulating the predictivity of invalid cues, which required reorienting to the target. Three differences in terms of spatiotemporal dynamics emerged, all in the direction of enhanced neural modulation for nonpredictive as compared to predictive invalid cues. First, nonpredictive invalid cues increased activation over ventral regions of the temporal and occipital lobes of the left hemisphere before the presentation of the target (likely the period in which attentional orienting is implemented). This is in line with findings from our research group, in which some attention-consciousness modulations were also left lateralized: for example, we observed left FEF activation in the interaction between orienting - and consciousness (Chica et al 2013b), and left IFG-ACC and left FEF-IPL pairwise connectivity on the executive attention-consciousness study (Martín-Signes et al 2019). Last, left parietal cortex was overall more active for seen over unseen trials (Chica et al 2016, Chica et al 2013b). Similarly, Chen et al (2019) showed robust structural connectivity patterns between the ventral surface of the occipito-temporal lobe and frontoparietal regions mediated by the attention-related Superior Longitudinal Fasciculi (see also Bartolomeo et al 2012, Chica et al 2018, Thiebaut De Schotten et al 2011). More recently, Sani and colleagues complemented this structural evidence with fMRI data from participants performing a high-load attentional task, and uncovered the existence of a ventro-temporal cortical node for object-based endogenous attention (Sani et al 2021). Our results seem to support this evidence regarding the existence of attention-related nodes in the inferior temporal cortex associated with attentional orienting.

Second, nonpredictive invalid cues increased activation in the temporo-occipital node and in the anterior insular cortex of the left hemisphere during the early stages of target processing (i.e., around 100ms after target presentation). The anterior insular cortex participates in attentional function (Xuan et al 2016) and cognitive control of information processing (Wu et al 2020a, Wu et al 2020b, Wu et al 2019). The temporal resolution of MEG recordings allows us to infer about the potential role of this frontal region when reorienting is necessary. Approximately 100ms following the presentation of an invalidly cued reported target, participants become aware of the mismatch between cue and target location. Doricchi et al (2010) showed that the brain deals with such discrepancy by recruiting frontal regions when the cue is invalid, and more posterior regions when the cue is valid. They proposed that target-related activity in the left TPJ encodes the matching between expected and actual target position in valid trials during stimulus-driven attention (similar to our experiment with nonpredictive cues) (but see Shulman et al 2007, Shulman et al 2003). Interestingly, however, Doricchi et al (2010) also observed enhanced activation of the IFG-MFG area in response to invalidly cued targets following predictive cues. Thus, the response of IFG-MFG to invalid cues was increased for predictive as compared to nonpredictive cues, because predictive cues induce strong focusing of endogenous attention on valid locations. Our results add to this knowledge by showing increased left-lateralized activation during attentional reorienting for nonpredictive over predictive invalid trials, further stressing the role of ventral frontal regions for template matching in stimulus-driven attention.

Third, nonpredictive invalid cues increased activation in temporo-parietal and temporo-occipital regions of the right hemisphere during later stages of target processing (starting approximately 250ms after target presentation). Target-related activity in the lateral occipital cortex and occipito-temporal system, bilaterally, has been previously associated with the report of seen near-threshold stimuli compared to unseen (Liu et al 2012). Our results confirm and specify these findings, because we observed a right-lateralized activation in temporo-parietal and occipito-parietal regions in the late stages of processing for seen targets as compared to unseen targets, and this activation was greater for nonpredictive than predictive cues. An alternative explanation for the increased activation of these clusters for seen over unseen targets when invalid nonpredictive cues are presented may be the existence of a *posterior hot zone* postulated by the Integrated Information Theory (Tononi et al 2016), as well as high-level visual nodes implicated in the Global Neuronal Workspace Theory (Mashour et al 2020). In line with this evidence, in a recent study (Liu et al 2022), we conducted intracerebral recordings in drug-refractory epilepsy patients undergoing this procedure for clinical reasons. Results showed a cluster of activity in electrodes located in the left-hemisphere prefrontal cortex and temporoparietal area showing activation to conscious reports that did not interact with attention. This result is consistent with the behavioral pattern showing increased percentage of detected targets when the invalid cue was nonpredictive rather than predictive, what we have defined here as the *cost* of “inappropriate” attentional orienting toward invalid predictive cues.

While adjudicating between rival theories of consciousness is beyond the scope of the current manuscript, our results elucidate how attention interacts with visual conscious processing. We observed greater reportability of target stimuli under conditions of increased attention, a finding in line with theories supporting that consciousness cannot occur without attention (the gateway hypothesis, Posner 1994) and pointing at the critical role of frontal regions in supporting conscious visual processing (the global neuronal workspace, Dehaene et al 2006, Mashour et al 2020), at least for near-threshold targets. Observing no effects of the cue validity on conscious perception, along with the absence of interaction between attention and consciousness in frontoparietal activations, would have favored the cumulative influence hypothesis (Tallon-Baudry 2012, Wyart & Tallon-Baudry 2008). Here, we show how valid predictive visual cues enhance visual conscious perception by recruiting frontoparietal regions, bilaterally. Here, we show how valid predictive visual cues facilitate visual conscious processing, pointing at one way in which attention and consciousness are related in the experimental setting used in this study. Behavioral data suggest that the most effective attentional cues to improve target perception are those predictive of the future location of the target, and occurring in close spatial proximity to it (Chica et al 2011a). Our results suggest plausible neural mechanisms for this effect.

A few comments and limitations need to be addressed. First, it remains possible that attention is required for conscious processing only when there is some competition between the stimuli to be resolved (Davidson et al 2018, Tsuchiya & Koch 2016), which was the case in our setting with two possible target locations. Alternatively, even an isolated stimulus might need some attentional capture to be consciously processed. Evidence from visual mental imagery studies, showing the implication of frontoparietal attention networks within the conscious imagination of an object in its absence, and thus without any competition (Spagna et al 2020) might support this possibility, which needs to be empirically assessed.

Two additional methodological aspects of our experiments deserve additional discussion: (1) cross-experiment comparisons are performed, even though the two participant samples only minimally overlapped (i.e., only two subjects performed both experiments). Therefore, a confounding factor may be at play, with the difference between nonpredictive and predictive cue actually being related to individual differences in our participants. While this option cannot be ruled out entirely in this study, the same analyses were conducted on the two overlapping participants, and led to results that resemble those observed at the group level – although not statistically significant due to the decreased power. (2) The number of catch (target-absent) trials differed between experiments with the nonpredictive cue experiment having a greater number of those (160 catch trials; 20% of the total) compared to the predictive cue experiment (112 catch trials; 14.5% of the total). This difference was due to the following reasons: (1) we wanted to the keep the total experiment duration similar across experiments (around 800 trials and 40 minutes), (2) while being mindful of the final number of trials that will undergo our analyses. Indeed, the *PredCue* (43%) and *NonPredCue* (40%) experiments differed minimally in the percent of consciously reported trials that underwent behavioral analyses after (a) excluding trials during pre-processing; (b) dropping about 50% due to the staircase procedure.

In conclusion, we show three patterns of interaction between attention and visual conscious processing that depend on the specific attentional component manipulated, contributing to the longstanding debate over theories of consciousness (recently discussed in Del Pin et al 2021, Michel et al 2019) and associated neural correlates (Chica et al 2016, Mashour et al 2020, Morales & Lau 2020, Raccah et al 2021). First, both predictive and nonpredictive valid visual cues modulated behavioral performance by increasing the number of detected targets. Predictive cues also modulated the response criterion and interacted with consciousness at the neural level, with invalid cues showing delayed frontoparietal involvement compared to valid cues, reflecting the cost of attentional reorienting. Third, cross task comparisons also showed that invalid predictive cues had the lowest percentage of detected targets and reduced neural activation compared to nonpredictive cues, a difference due to the probabilistic expectations embedded in the cue predictivity. Altogether, the comparison of the spatiotemporal dynamics underlying the interaction between nonpredictive and predictive attention with consciousness shown here confirmed that these distinct contributions can be observed both on behavioral and on neural measurements. Our findings underline the role of the frontoparietal networks in supporting the interaction between attention and consciousness, and specify occipito-temporal, anterior insular, and parieto-temporal markers associated with the conscious processing of invalidly cued targets, which may counter the cue-induced distraction for targets occurring at the uncued location.

## Supporting information

Supplementary Method and Results

## Author Contribution

D.B., A.B.C., and P.B. designed the experiments; A.S., D.B., Z.R., A.B. C. analyzed the data. A.S., A.B.C. and P.B. draft the report. All authors discussed the results and revised the report.

## Acknowledgment

We thank Fabrizio Doricchi, Isabella Elaine Rosario, and Catherine Tallon-Baudry for providing extensive comments on this manuscript.

## Data Availability

Behavioral data and the code to reproduce the figures (built using Python on Spyder) can be found on the <u>GitHub page of A.S.</u>

