## Supplementary Table 1 for "The cost of attentional reorienting on conscious visual perception: an MEG study"

| **PerPred: n trials remaining after artifact rejection** | | | | | | | | | | | | | |
| --- | --- | --- | --- | --- | --- | --- | --- | --- | --- | --- | --- | --- | --- |
| subject | SRI | SRV | SLI | SLV | URI | URV | ULI | ULV |  | Sum | Sum of Valid | Sum of Invalid | Difference V - I |
| 1 | 20 | 92 | 32 | 84 | 45 | 57 | 32 | 68 |  | 430 | 301 | 109 | 192 |
| 2 | 24 | 85 | 39 | 85 | 61 | 110 | 55 | 107 |  | 566 | 387 | 155 | 232 |
| 3 | 35 | 105 | 41 | 96 | 48 | 66 | 33 | 85 |  | 509 | 352 | 122 | 230 |
| 4 | 42 | 85 | 30 | 80 | 46 | 77 | 46 | 74 |  | 480 | 316 | 122 | 194 |
| 5 | 41 | 135 | 50 | 104 | 50 | 54 | 52 | 95 |  | 581 | 388 | 152 | 236 |
| 7 | 32 | 85 | 37 | 93 | 55 | 97 | 50 | 85 |  | 534 | 360 | 142 | 218 |
| 8 | 31 | 145 | 70 | 92 | 62 | 57 | 33 | 111 |  | 601 | 405 | 165 | 240 |
| 9 | 25 | 153 | 62 | 83 | 66 | 35 | 34 | 103 |  | 561 | 374 | 162 | 212 |
| 11 | 12 | 153 | 61 | 27 | 78 | 35 | 35 | 157 |  | 558 | 372 | 174 | 198 |
| 12 | 34 | 121 | 64 | 79 | 54 | 59 | 29 | 107 |  | 547 | 366 | 147 | 219 |
| 13 | 16 | 152 | 48 | 73 | 80 | 36 | 35 | 114 |  | 554 | 375 | 163 | 212 |
| 14 | 29 | 103 | 48 | 113 | 55 | 64 | 46 | 74 |  | 532 | 354 | 149 | 205 |
| 15 | 21 | 114 | 44 | 64 | 65 | 72 | 42 | 109 |  | 531 | 359 | 151 | 208 |
| 17 | 34 | 60 | 13 | 71 | 47 | 101 | 54 | 90 |  | 470 | 322 | 114 | 208 |
| Sum | 396 | 1588 | 639 | 1144 | 812 | 920 | 576 | 1379 |  | 7454 | 5031 | 2027 | 3004 |
| **PerNonPred: n trials remaining after artifact rejection** | | | | | | | | | | | | | |
| subject | SRI | SRV | SLI | SLV | URI | URV | ULI | ULV |  | Sum | Sum of Valid | Sum of Invalid | Difference V -I |
| 1 | 84 | 76 | 87 | 110 | 56 | 68 | 52 | 29 |  | 562 | 283 | 195 | 88 |
| 2 | 93 | 51 | 44 | 90 | 37 | 85 | 82 | 46 |  | 528 | 272 | 163 | 109 |
| 3 | 60 | 68 | 63 | 70 | 44 | 41 | 42 | 36 |  | 424 | 215 | 149 | 66 |
| 4 | 96 | 27 | 15 | 95 | 20 | 81 | 100 | 22 |  | 456 | 225 | 135 | 90 |
| 5 | 55 | 105 | 99 | 55 | 72 | 30 | 33 | 73 |  | 522 | 263 | 204 | 59 |
| 6 | 26 | 85 | 74 | 24 | 56 | 26 | 22 | 60 |  | 373 | 195 | 152 | 43 |
| 7 | 45 | 64 | 62 | 79 | 74 | 64 | 75 | 47 |  | 510 | 254 | 211 | 43 |
| 8 | 61 | 84 | 74 | 50 | 72 | 46 | 49 | 80 |  | 516 | 260 | 195 | 65 |
| 9 | 77 | 50 | 54 | 85 | 35 | 60 | 62 | 35 |  | 458 | 230 | 151 | 79 |
| 10 | 54 | 103 | 97 | 60 | 79 | 42 | 41 | 71 |  | 547 | 276 | 217 | 59 |
| 11 | 57 | 65 | 72 | 64 | 76 | 58 | 53 | 62 |  | 507 | 249 | 201 | 48 |
| 12 | 42 | 99 | 108 | 52 | 65 | 11 | 14 | 56 |  | 447 | 218 | 187 | 31 |
| 13 | 50 | 89 | 62 | 80 | 66 | 31 | 55 | 36 |  | 469 | 236 | 183 | 53 |
| 14 | 88 | 55 | 53 | 74 | 38 | 59 | 70 | 48 |  | 485 | 236 | 161 | 75 |
| 15 | 24 | 63 | 54 | 31 | 37 | 22 | 27 | 45 |  | 303 | 161 | 118 | 43 |
| 16 | 48 | 59 | 55 | 45 | 62 | 57 | 59 | 72 |  | 457 | 233 | 176 | 57 |
| 17 | 52 | 51 | 53 | 75 | 72 | 73 | 74 | 45 |  | 495 | 244 | 199 | 45 |
| 18 | 21 | 90 | 65 | 37 | 95 | 28 | 46 | 76 |  | 458 | 231 | 206 | 25 |
| Sum | 1033 | 1284 | 1191 | 1176 | 1056 | 882 | 956 | 939 |  | 8517 | 4281 | 3203 | 725 |

Note: S = seen; U = unseen; R = right; L = Left; I = Invalid; V = Valid.
