## Supplementary Method and Results for "The cost of attentional reorienting on conscious visual perception: an MEG study"

**Supplementary Results**

A similar ANOVA conducted on RTs revealed a main effect of *Consciousness* (F_(1,17)_ = 102.65; *p* < .0001; η^2^ = .86), because participants were slower for *Seen* targets (1034 ± 115ms) than for *Unseen* targets (601 ± 190ms). No other factors or interactions reached significance.

***Does the validity of nonpredictive cues affect neural responses associated with the conscious report of visual Gabor targets?*** We addressed this question by examining brain responses to seen and unseen targets separately for valid and invalid nonpredictive cues. Source analysis of the MEG signal revealed that conscious perception was associated with right-lateralized frontoparietal feed-forward and feedback sweeps. Two clusters exceeding the threshold of randomization distribution under H0 emerged for the seen *vs* unseen comparison (both *p*s < 0.001) in the time window of 400 - 800ms after cue onset. The first cluster was in the right hemisphere, the second in the left hemisphere. Both clusters started in the occipital cortex and afterwards extended to the frontoparietal network and temporal regions, bilaterally (see **Fig S1**). The differences for the main effects of *Validity* (valid, invalid), for the main effect of *Visual Field* (left, right), and for the interactions did not reach statistical significance. Control analyses showed that there was no significant difference between the number of MEG trials for left- and right-sided targets (left visual field, mean ± SD: 235.39 ± 32.49; right visual field, 235.5 ± 30.02; Wilcoxon signed-rank test = 85; *p* = 1; Bayesian Wilcoxon signed-rank test BF_10_ = 0.57, with median posterior *δ* = -0.05, 95% CI [-1.09, 0.96]).

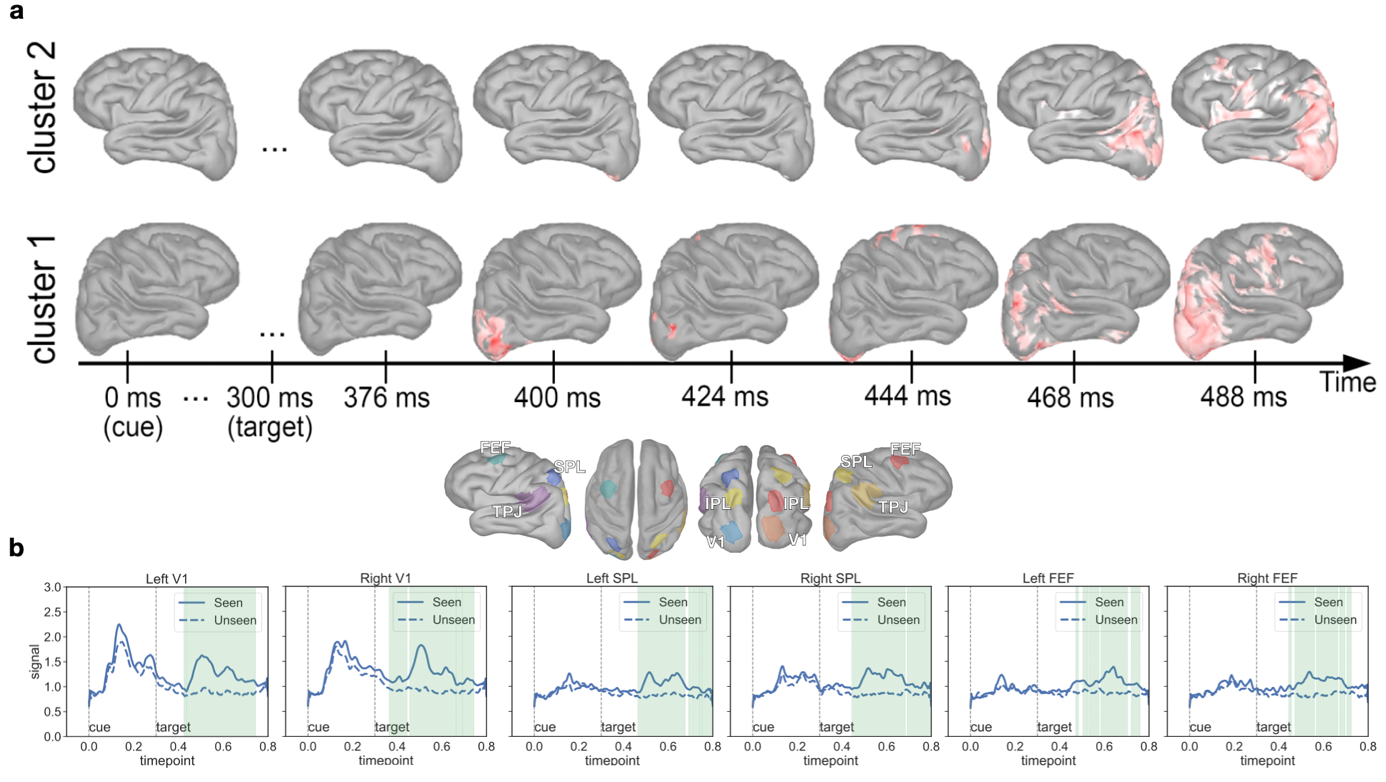

**Figure S1.** **a.** When preceded by nonpredictive cues, seen targets evoked two clusters of brain activity compared to unseen targets. Cluster 1 occurred in the 400 - 800ms time window after cue onset, was lateralized to the right hemisphere, and encompassed a frontoparietal *feed-forward* and *feedback sweeps* (around 456ms), with subsequent diffusion to widespread bilateral activation. Cluster 2 occurred also in the 400 - 800ms time window after cue onset, but was lateralized to the left hemisphere, and encompassed a widespread brain activation. **b.** Average signal changes in the ROIs separately for the Seen (solid line) and Unseen (dashed line) condition. The area in green highlights the time interval in which cluster-corrected analysis showed a significant difference between the two signals.
